# Proteomics-Enhanced AI-Digital Pathology in Metastatic Mucinous Colorectal Carcinoma: A Case Report

**DOI:** 10.64898/2026.03.02.709044

**Authors:** Livia Fülöp, Balázs Szigeti, Jéssica Guedes, Nicole Woldmar, Henriett Oskolás, Matilda Marko-Varga, Roger Appelqvist, Elisabet Wieslander, Krysztof Pawlowski, Leticia Szadai, Lukas Christersson, Johan Malm, István Balázs Németh, Marcell A. Szasz, Jeovanis Gil, György Marko-Varga

**Affiliations:** National Koranyi Institute of Pulmonology, Budapest, Hungary; Section for Clinical Chemistry, Department of Translational Medicine, Lund University, Lund, Sweden; Clinical Protein Science & Imaging, Biomedical Centre, Department of Biomedical Engineering, Lund University, Lund, Sweden; Department of Molecular Biology, University of Texas Southwestern Medical Center, Dallas, Texas, USA; Department of Dermatology and Allergology, University of Szeged, Szeged, Hungary; Department of Thoracic Surgery, Semmelweis University and National Institute of Oncology, Budapest, Hungary; Chemical Genomics Global Research Lab, Department of Biotechnology, College of Life Science and Biotechnology, Yonsei University, Seoul, Republic of Korea; 1st Department of Surgery, Tokyo Medical University, Tokyo, Japan

**Keywords:** Colon cancer, proteomics, subtypes, stratification, histopathology, tumor microenvironment, metastases

## Abstract

Mucinous colorectal carcinoma (CRC) is a distinct histomorphological subtype characterized by abundant extracellular mucin that may promote immune evasion and chemoresistance. We describe a metastatic mucinous CRC case integrating digital pathology and proteomics to investigate disease progression and therapy resistance. Formalin-fixed paraffin-embedded samples from the primary tumor, peritoneal metastasis, and hepatoduodenal ligament metastasis of a 56-year-old patient were analyzed. Whole-slide imaging with QuPath-based AI enabled detailed histological annotation, while data-independent acquisition mass spectrometry identified over 6,000 proteins. Digital pathology revealed extensive mucin pools, architectural evolution from heterogeneous glandular patterns in the primary tumor to cribriform morphology in advanced metastases, and immune cell exclusion from mucin-rich regions. Proteomics revealed metabolic reprogramming, suppressed antigen presentation, and stage-specific activation of inflammatory, angiogenic, EMT, and PI3K/AKT/mTOR–MYC signaling pathways, consistent with proliferative and therapy-resistant phenotypes.Integration of AI-assisted histopathology with spatial proteomics highlighted the mucin barrier as a key mediator of immune evasion and chemoresistance. These findings support a personalized therapeutic framework targeting mucin-associated mechanisms alongside pathway-directed inhibitors, suggesting that spatial multi-omics may guide precision management strategies for aggressive mucinous colorectal cancer.

## 1. Introduction

Today, colorectal cancer (CRC) remains a major global health burden, with an estimated 1.85 million new cases reported in 2018 and continuing to rise in many regions (1). It ranks as the second most common cancer in women and the third in men, highlighting its broad epidemiological and clinical significance (2). The development of CRC is strongly associated with Western dietary patterns (2). These risk factors are particularly prevalent in high-income countries, where CRC incidence historically has been the highest (3). While recent data indicate a stabilization or even a decline in incidence among older adults in several developed nations attributable in part to improved screening and early detection programs a disturbing countertrend has emerged: a consistent and unexplained rise in CRC incidence among young adults under 50 years of age, particularly in countries such as Australia, Canada, the United States, and parts of Europe (4).

This shift in age-related incidence is of great concern, as early-onset CRC often presents at a more advanced stage, with aggressive histological features, and is more likely to be misdiagnosed or delayed in diagnosis due to the low clinical suspicion in younger populations. It underscores the urgent need to revisit current screening guidelines and to develop age-tailored strategies for prevention and early intervention. Histologically, the vast majority of CRCs, which can be defined by at least submucosal level of colonic wall invasion, are classified as adenocarcinomas. Among these, mucinous adenocarcinoma (MAC) represents a distinct histomorphological subtype, characterized by large extracellular mucin pools occupying more than 50% of the tumor volume (5). MAC occurs in approximately 5–20% of CRC cases and is often associated with specific clinical features, including younger age at onset, proximal colon location, and resistance to certain chemotherapies (5). Its molecular profile frequently overlaps with microsatellite instability (MSI) and CpG island methylator phenotype (CIMP), further emphasizing its distinct biology (5). In light of the evolving landscape of CRC, improved understanding of tumor subtypes, patient demographics, and molecular underpinnings is essential. As for recent advances in molecular classification systems, key efforts include refining consensus molecular subtypes (CMS), elucidating the role of the tumor microenvironment in immune evasion and treatment response, and leveraging transcriptomic signatures for patient-specific clinical decision-making (6). Previous studies (7,8) have updated the molecular pathological classification of CRC, while a large-scale genomic analysis of 2,023 CRC tumors provided detailed insights into mutation spectra, pathway alterations, and potential drug targets (9). In parallel, recent immunopathological investigations (10) highlight the prognostic significance of the Immunoscore across CMS classes, supporting its integration into routine pathological evaluation. In addition, recent advances in artificial intelligence (AI) and digital pathology are increasingly enabling the integration of complex multimodal datasets, including histomorphology, proteomics, and molecular signatures (11,12). By combining computational image analysis with proteomic profiling, emerging frameworks aim to refine tumor characterization and support precision oncology strategies. Together, these developments underscore the critical importance of combining morphological, molecular, and immunological data to guide the next generation of diagnostic and therapeutic strategies for CRC.

## 2. Methods

### 2.1. Case presentation

The patient is a 58-year-old man who initially presented in 2019 with a sessile adenomatous polyp, identified by colonoscopy, in the sigmoid colon, which upon resection the initial histopathological report revealed invasive adenocarcinoma with mucinous features. A subtotal colectomy was performed, including the site of the previously resected sessile lesion, and the pathologic report was signed out as a pT3N1 mucinous adenocarcinoma. Despite adjuvant chemotherapy (capecitabine and oxaliplatin), the disease recurred. Over the next four years, the patient underwent multiple surgeries for metastatic relapses, including cytoreductive surgery with hyperthermic intraperitoneal chemotherapy (HIPEC) in 2021 for peritoneal metastases. In 2023, new metastases appeared along the peritoneum and in the hepatoduodenal ligament (near the porta hepatis), confirmed by MRI. The patient also developed a metachronous prostate adenocarcinoma during this period, which was managed with hormonal therapy and radiotherapy, achieving control independent of the CRC. In June 2024, a second cytoreductive surgery was undertaken to remove extensive peritoneal tumor burden, including implants on the abdominal wall and within subcutaneous tissue. By this time, the peritoneal disease showed a gelatinous, mucin-rich appearance consistent with pseudomyxoma peritonei-like spread. The patient’s clinical trajectory and treatments are summarized in **Figure 1**.

**Figure 1.**
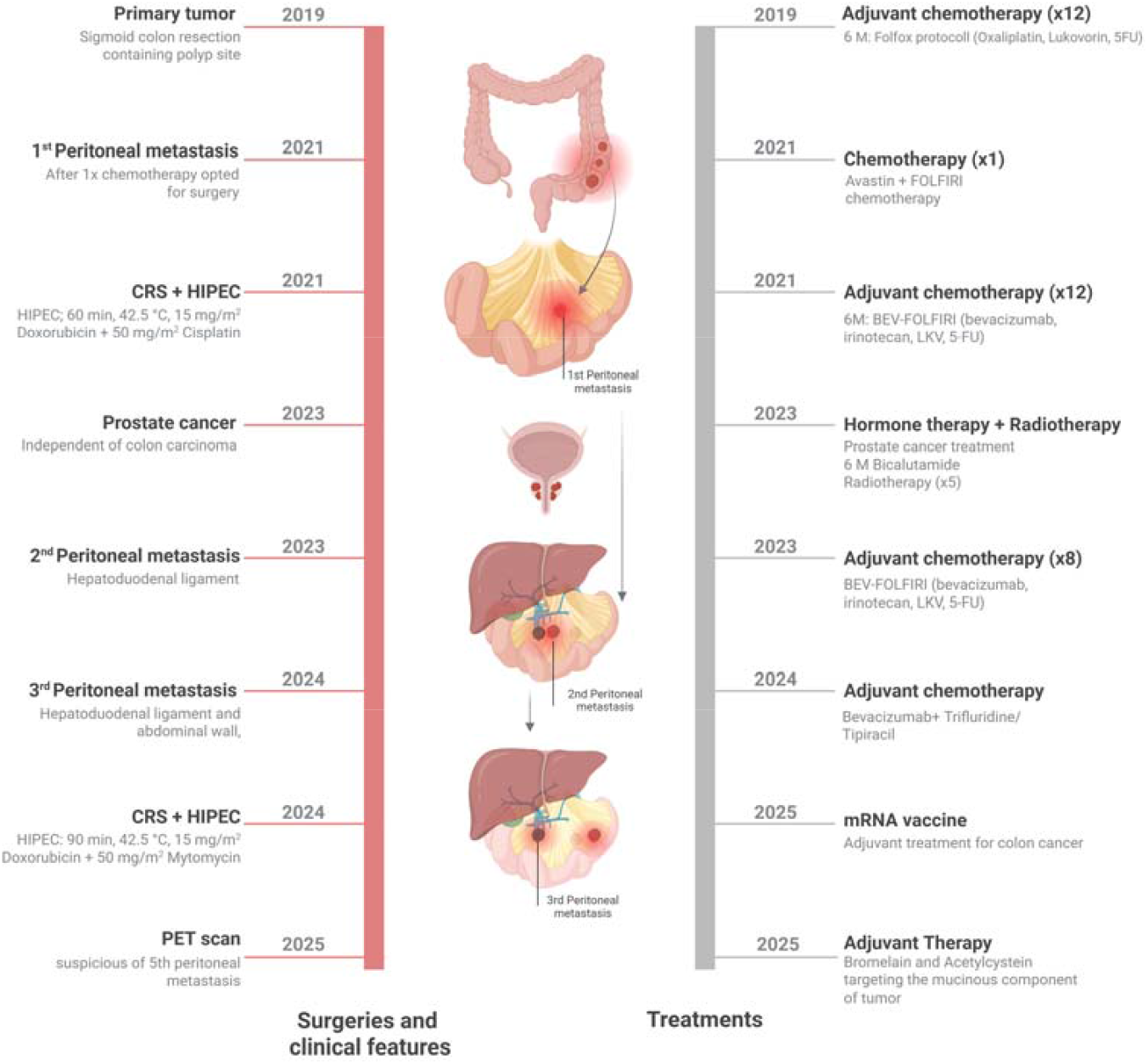
Timeline of the patient’s disease course and interventions. Key milestones include the initial diagnosis of a mucinous sigmoid colon carcinoma in 2019, surgical resections of the primary tumor, the onset of peritoneal metastases in 2021 requiring cytoreductive surgery with HIPEC, a second malignancy (prostate cancer) treated in parallel, and further metastatic progression to the hepatoduodenal ligament and abdominal wall by 2023–2024. Therapeutic interventions (surgery, chemotherapy, radiotherapy, and HIPEC) and their timing are indicated, illustrating the aggressive and recurrent nature of the disease.

### 2.2. Patient samples

The study was carried out in strict accordance with the Declarations of Helsinki and was approved by the national-level Ethics Committee (Hungarian Scientific and Research Ethics Committee of the Medical Research Council, BM/15061-1/2023). Formalin-Fixed Paraffin-Embedded tissue (FFPE) tissue samples from the primary tumors of patients diagnosed with metastatic prostate cancer were obtained from the Buda Hospital of the Hospitaller Order of Saint John of God (Budapest, Hungary).

All resected tumor specimens were FFPE for histopathological analysis. Hematoxylin and eosin (H&E) stained whole-slide images were digitized at 40× equivalent resolution. Board-certified pathologists reviewed the slides to identify regions of interest, including varying tumor architectures, mucin pools, invasion fronts, and host tissue interfaces. We employed QuPath (v0.4.x), an open-source digital pathology platform (13), to assist in annotation and analysis. Custom Groovy scripts were developed to extend QuPath’s functionality, enabling efficient management of complex annotations and extraction of quantitative features for downstream correlation with proteomic data. For example, a *“c2e_v2.groovy”* script was used to export annotation metrics (areas of tumor vs. mucin vs. stroma, etc.) as structured data, facilitating direct comparison with proteomic abundances. These tools enhanced the reproducibility of our spatial analysis by tracking annotator inputs and ensuring consistency across images. Upon histomorphological inspection throughout the patient’s specimens, we microscopically noted a predominantly immune-excluded pattern: lymphocytes, when present, were confined to stromal rims around tumor/mucin pools, with virtually no penetration into tumor islets.

### 2.3. Proteomic analysis

FFPE blocks corresponding to the pathologist-reviewed specimens were processed as whole tissue sections (WTS). Samples were processed as previously described (14–16). Briefly, sections were deparaffinized, proteins extracted in 5% SDS/100 mM TEAB (pH 8.0) with 25 mM DTT at high temperature, and lysates sonicated to enhance solubilization. After quantification, cysteines were alkylated (IAA), and digestion was performed using S-Trap devices, followed by trypsin at 1:25 (enzyme:protein). Peptides were eluted stepwise, dried, and reconstituted in 0.1% TFA/2% ACN for LC-MS analysis.

Peptide mixtures (1 µg) were analyzed by nanoLC–MS/MS on an Ultimate 3000 coupled to a Q-Exactive HF-X. Samples were trapped and separated on a 50 cm C18 column with a 100-min non-linear gradient at 300 nL/min and 60 °C. Data were acquired in variable-window DIA to maximize coverage and quantification consistency. Run quality was monitored via identification rates, TIC stability, mass accuracy and retention-time behavior (17).

Raw DIA files were processed in DIA-NN (directDIA) against the UniProt human reference proteome, controlling 1% FDR at peptide and protein levels. Protein group intensities were log□-transformed and median-normalized; basic filtering removed features with excessive missingness. Differential abundance focused on prespecified contrasts aligned to the clinical trajectory (primary vs adjacent colon; peritoneal metastasis vs adjacent colon; hepatoduodenal metastasis vs peritoneal metastasis). Functional interpretation used GSEA to evaluate processes highlighted by pathology, e.g., extracellular matrix/mucin, antigen processing/presentation, EMT, angiogenesis, IL-6/JAK/STAT3, PI3K/AKT/mTOR, MYC targets, oxidative phosphorylation and fatty-acid metabolism. Integration with histology was performed at the specimen level.

## 3. RESULTS

### 3.1. Primary Tumor Analysis (Sigmoid Colon) – Histology and Proteomic Signature

The primary tumor presented in **Figure 2** originates from a conventional tubular type of sigmoid colon polyp and has been identified as a colorectal adenocarcinoma with mucinous features. Histologically, this subtype of tumor, which are typically observed in 5–10% of colorectal adenocarcinoma cases, displays glandular epithelial formations floating within abundant mucinous substance. These mucinous components are associated with poorer systemic treatment responses, particularly in metastatic settings, where drug penetration and immune engagement are often compromised. Despite their clinical significance, no comprehensive studies to date have systematically explored the molecular pathology underlying this heterogeneous mucinous phenotype in solid tumors. Our integrated spatial proteomics and histopathological analyses reveal the presence of distinct epithelial phenotypes within the tumor mass, characterized by differential protein expression and architectural segregation. Notably, we observed a consistent correlation between mucin-rich regions and the epithelial compartment, suggesting a functional interaction that may influence tumor plasticity, immune evasion, and therapeutic resistance.

**Figure 2.**
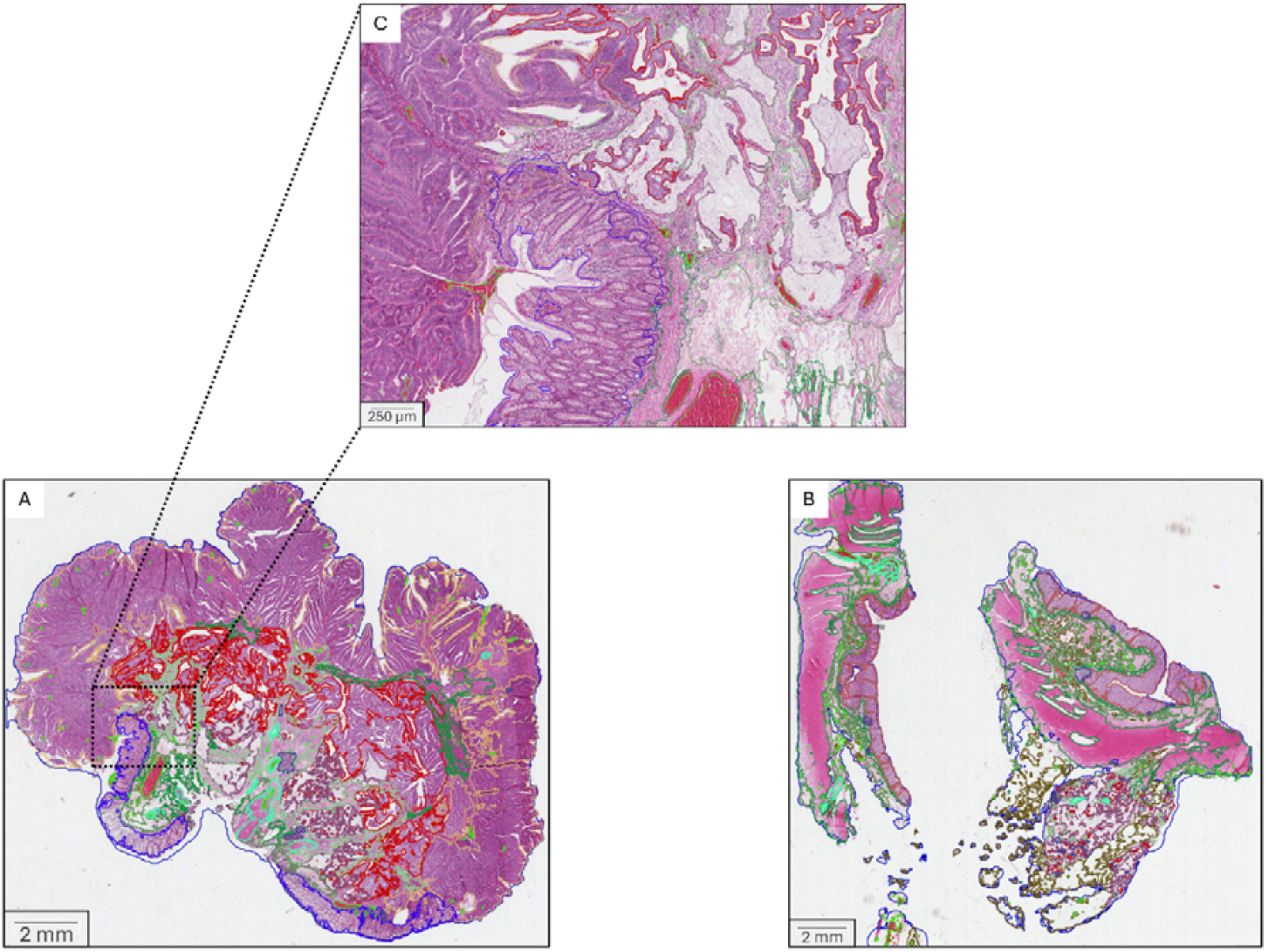
Histological images generated from the primary colon tumor **(A)**. Illustrated examples of the QuPath annotation procedure developed and implemented throughout the study, shown here using primary tumor tissue as a representative compartment captured from the whole-slide image. The left panel **(A)** originates from a conventional sigmoid polyp and demonstrates four distinct phenotypic characteristics within the tissue microarchitecture. The right panel **(B)** shows the sigmoid resection specimen containing the primary tumor site. The sectioned plane reveals remnants of invasive tumor predominantly located within the subserosal adipose tissue beneath the muscularis propria layer **(C)**. High-magnification view of the highlighted region from image **(A)**, providing a detailed zoom-in **(C)** of the annotated area. Sampled regions are marked with blue outlines. Tumor cell clusters or tumor tissue in red, adipose tissue in brown, blood vessels or stromal components in light green. Dark green denotes connective tissue or smooth muscle in dark green, purple indicates mucin, and lymphoid aggregates are outlined in dark blue. Normal colonic mucosa is annotated in blue **(A)**, whereas normal colonic epithelium in orange **(B)**. Scale bars: **(A)** 2 mm; **(B)** 2 mm; **(C)** 250 µm.

In order to support high-throughput and reproducible histopathological image analysis, we optimized the QuPath software platform (v0.4.x) by integrating a suite of custom Groovy scripts specifically designed to streamline annotation workflows. In addition, a major goal within the study was to make an overall improvement on the quantitative data extraction. The buttons groovy script introduces intuitive on-screen UI components that display annotator identity and contact details, this is of mandatory importance, that enhances the traceability as well as the collaborative communication within pathology departments across multiple hospitals. c2e_v2.groovy enables extraction of quantitative metrics, such as the annotation area, the intensity values, and the subclass identifiers, we exported in *.csv formats and executed downstream statistical evaluation, interfacing the proteomic correlation. To ensure the quality and performance stability throughout the batch processing, the checker.groovy monitors the RAM usage in real-time. This is a practical approach that helps mitigate system crashes and processing delays. Earser.groovy facilitates bulk deletion of redundant or erroneous annotations, we found this to be very helpful as we were annotating across our team interactions, that helps preserving the dataset accuracy and consistency. Finally, subclassifier_v4.groovy will calculate and export aggregated the area metrics (µm^2^) and the region-specific ratios for annotated tumor compartments and phenotypes (see below, A–E). With this feature, we are supporting a detailed morphometric analysis. Altogether, these tools are scalable and form a robust infrastructure for integration with advanced AI-based morphological analysis including the multi-omics data.

The annotation colors used throughout this study correspond to specific tissue types. Sampled regions are delineated by blue or green outline. Tumor cell clusters or tumor tissue are indicated in red, adipose tissue in brown, and blood vessels or stromal components in light green. Dark green denotes connective tissue or smooth muscle, purple indicates mucin, and lymphoid aggregates are outlined in dark blue. In **Figure 2A**, normal colonic mucosa is annotated in blue, whereas in **Figure 2B**, normal colonic epithelium is annotated in orange. Comprehensive histopathological evaluation of the surgically resected tumor specimens revealed five distinct phenotypic patterns of malignant epithelial organization. These patterns demonstrated varying degrees of architectural complexity, nuclear pleomorphism, and levels of histopathological differentiation, ranging from well, organoid looking to lesser differentiated, high grade patterns. Collectively, they appear to represent a continuum of spatial and temporal progression stages in epithelial tumor evolution, from early localized lesions to more aggressive, metastaszing phenotypes. Each of these histological profiles was associated at the proteomics level with different interactions towards the surrounding stroma, including immune infiltration and desmoplastic remodeling, suggesting a dynamic crosstalk between malignant cells and their microenvironment throughout tumor progression. Furthermore, these histopathological phenotypes were not singularly represented in each specimen, but variously mixed components of malignant epithelia were present in each manifestation and throughout tumor progression.

Based on the in-depth digital pathological examination of the malignant epithelia, the phenotypes were characterized as follows:

**A)** Dysplastic Epithelial Phenotype “Invasive elements arising from Adenomatous polyp”: This phenotype is characterized by epithelial cells showing high grade dysplastic features, forming elongated crypts and glandular structures, marked with loss of nuclear polarity, marked cytological atypia and frequent mitoses. These structures appear as the “invasive component” within the primary polyp, infiltrating beyond the muscularis mucosa layer with malignant epithelia resembling the high-grade dysplastic part of the adenoma. This component represents a transition from high grade, in situ dysplasia to invasive carcinoma, suggesting a critical zone of early malignant transformation. Upon observation, this morphological appearance is only detectable within the primary setting.
**B)** Primitive Gland-Forming Phenotype: Composed of smaller epithelial clusters exhibiting early glandular differentiation, this phenotype is notable for the formation of rudimentary luminal spaces. The architecture resembles poorly developed glands or tubules, often arranged in variably cohesive nests. The cells display intermediate atypia, with focal polarity and partial preservation of glandular architecture, reflecting a relatively well differentiated infiltrative growth pattern. This phenotype may correspond to an early invasive adenocarcinoma component.
**C)** Epithelial Clumps and Variably Stratified Layers: This variant features cohesive malignant epithelial cells arranged in compact clumps or linear layers without clear glandular organization. The cellular arrangement is less organized, the malignant epithelia appear to be less differentiated. This suggests a degree of loss regarding epithelial cohesion, potentially reflecting epithelial-to-mesenchymal transition (EMT)-associated changes and increased migratory potential. Cytological atypia varies within these lesser differentiated epithelial formations, but mitotic activity remains relatively elevated.
**D)** Floating Individual Tumor Cells: The most dispersed phenotype, characterized by single malignant epithelial cells scattered throughout the tumor microenvironment represented by mucinous pools and scarce stromal elements. These isolated cells exhibit pronounced pleomorphism and high nuclear-cytoplasmic ratios, often with tendency of detachment from primary clusters. This pattern is commonly associated with advanced invasive behavior, increased metastatic potential, and poor prognostic outcomes, particularly in mucinous or signet-ring cell carcinoma subtypes.
**E)** Complex glandular/cribriform epithelial formations: The last type of epithelial morphologies we observed is characterized by less defined multiple glandular lumens containing large cohesive clusters of malignant cells. This phenotype is represented by relatively large epithelial formations floating in EC mucin, containing multiple (cribriform) lumens inside large glandular spearing cohesive cell groups. In addition, less defined appearances of these larger formations can be observed, represented by poorly differentiated cellular patterns. This phenotype mainly appeared in late metastatic settings.

By examining the primary tumor compartment in colon cancer, we successfully distinguished healthy tissue regions from those infiltrated by tumor cells using image-based analysis. Notably, the tumor-generated mucin aggregates exhibited altered structural and biochemical properties compared to control healthy internal mucin. These aberrant mucin forms create a dense barrier that significantly may hinder lymphocyte infiltration and impede effective drug delivery. This protective microenvironment contributes to immune evasion and limits therapeutic efficacy. Understanding these mucin-associated changes is essential for developing strategies to overcome treatment resistance in mucinous colon tumors (**Figure 3**)

**Figure 3.**
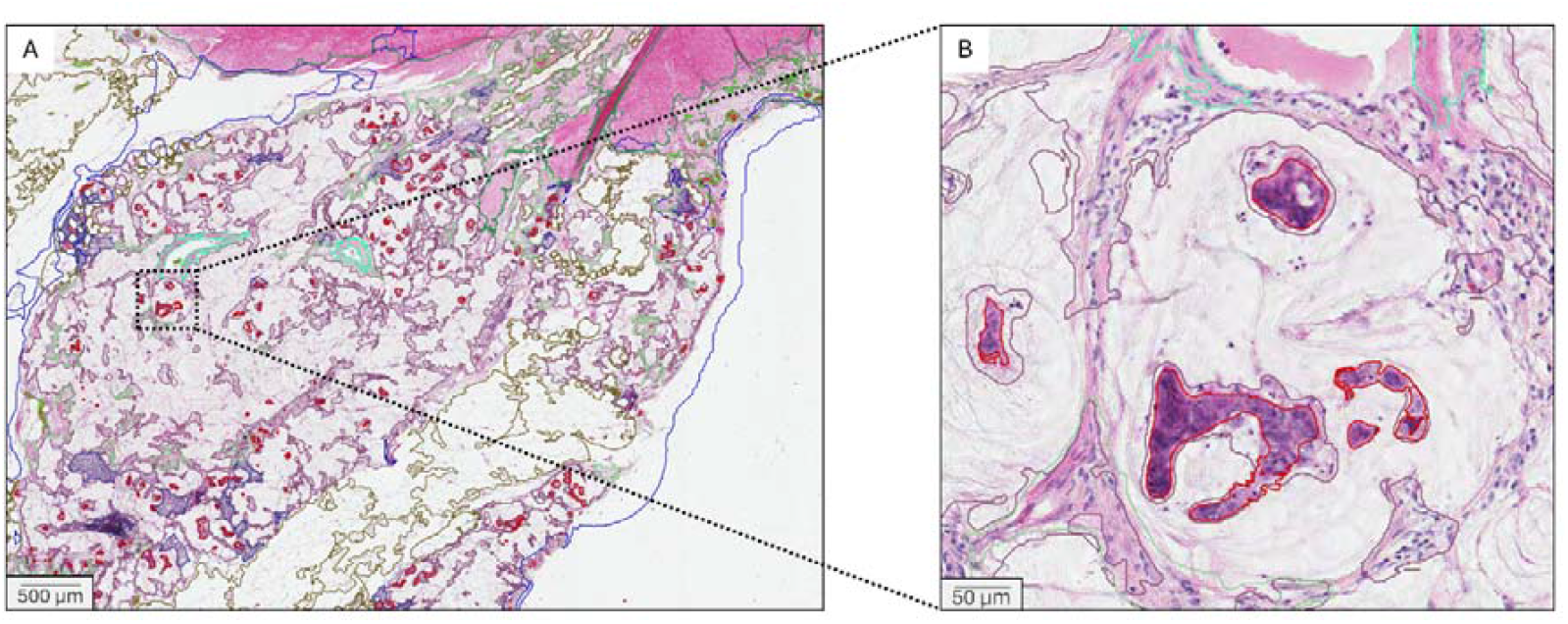
Histological images at varying magnifications obtained from primary colon tumor tissue of the resected sigmoid colon containing the cancerous polyp site. The left image **(A)** shows the muscularis propria layer in the upper region, beneath which subserosal adipose tissue of the colon is heavily infiltrated by mucinous adenocarcinoma cells (MAC). The right image **(B)** presents a higher-magnification view of the highlighted area from **(A)**, illustrating primitive gland formation, layered and scattered tumor cells, increased nuclear atypia, and loss of mucin integrity. These alterations are accompanied by changes in both structural and biochemical tissue properties, including aberrant gland formation and stromal remodeling. The visual contrast emphasizes the morphological and functional transformation associated with tumor progression in the mucinous subtype of colorectal cancer. Sampled regions are outlined by blue color. Tumor tissue is in red, adipose tissue in brown, blood vessels or stromal components in light green. Dark green denotes connective tissue or smooth muscle in dark green, purple indicates mucin, and lymphoid aggregates are outlined in dark blue. Scale bars: **(A)** 500 µm; **(B)** 50 µm.

### 3.2. Proteomic findings - Primary Tumor

The whole-tissue proteome of the primary sigmoid lesion mirrored its mucinous histopathology and immune-excluded microenvironment (**Figure 4**). MUC2, the dominant secreted mucin of colonic goblet cells, was overexpressed, consistent with the extensive extracellular mucin seen on H&E. In contrast, proteins underpinning adaptive immunity were broadly suppressed: elements of the antigen-processing/presentation machinery were reduced, interferon-stimulated programs and T-cell-attracting chemokines were under-represented, and complement/coagulation cascades were diminished. Taken together with the paucity of intratumoral TILs on digital pathology, these data support an immune-cold phenotype maintained by both a physical mucin barrier and tumor-intrinsic immune evasion (18). Functionally, the primary tumor showed a stress-adapted, cell-cycle-driven state with low metabolic tone. Hallmark/Reactome enrichment highlighted activation of the unfolded protein response, DNA repair, and proliferative axes, consistent with high replication pressure and proteostasis demand. In contrast, metabolic and inflammatory pathways were negatively enriched, including oxidative phosphorylation, fatty-acid metabolism, xenobiotic metabolism, reactive-oxygen-species pathway, heme metabolism, and adipogenesis, alongside hypoxia, angiogenesis, epithelial–mesenchymal transition, apical junctions, and multiple immune pathways. Of note, both KRAS signaling (UP/DN) Hallmark sets were negatively enriched in this specimen. Overall, the proteome depicts a mucin-rich, immune-excluded tumor that prioritizes cell-cycle progression and stress-response circuitry while down-tuning metabolic and immune/inflammatory activity, aligning with the histological phenotype and helping explain chemoresistance behind a dense mucin barrier.

**Figure 4.**
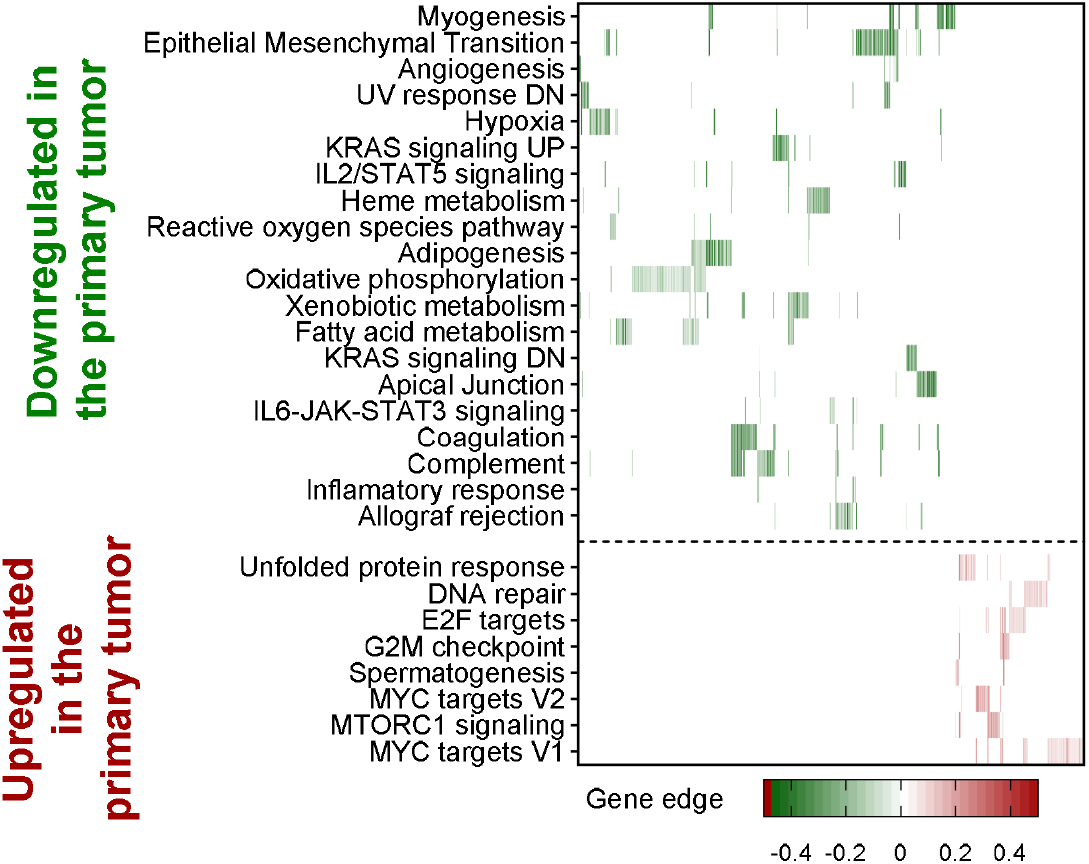
Differential protein regulation across signaling pathways and functional categories in colon tumor samples. The heatmap illustrates upregulated (red) and downregulated (green) proteins, grouped by biological function, including immune response, lipid metabolism, complement activation, and cell adhesion. Proteins involved in tumor-promoting pathways such as KRAS/JAK/STAT3, and innate immunity show significant upregulation in patient samples, while apolipoproteins and cholesterol metabolism proteins are predominantly downregulated. This functional stratification highlights distinct proteomic shifts associated with disease progression and potential response to therapy. Data represents fold changes relative to healthy controls, with statistical significance annotated.

### 3.3. Metastatic Tumor Tissue Presentation

Over a four-year clinical follow-up, the patient exhibited three separate relapse events, each exhibiting distinct pathological features. Through comprehensive histopathological and molecular profiling, we successfully identified three out of the four recognized phenotypic presentations associated with the patient’s primary mucinous adenocarcinoma. Notably, the dysplastic epithelial subtype was the only phenotype not observed across the multiple recurrence events. However, the presence of primitive gland forming, clump and layer forming and individually dispersed tumor cell patterns, were clearly confirmed through histological and molecular analysis and validated using transcriptomic data, reflecting the tumor’s capacity for phenotypic plasticity and intratumoral heterogeneity. These findings emphasize the importance of multi-sample, longitudinal analysis in capturing the dynamic evolution of tumor phenotypes, with implications for personalized treatment and resistance mechanisms.

Notably, within the metastatic relapses documented in May 2024, we identified the 5^th^, distinct complex glandular cribriform phenotype, which was scarcely present at earlier manifestations. This phenotype was localized within defined metastatic sites like the peritoneum and abdominal wall and presents unique morphological traits that distinguish it from early metastatic forms. The identification of this cribriform pattern may suggest a previously underrecognized route of mucinous colorectal adenocarcinoma progression or adaptive phenotype shift during metastatic dissemination. Ongoing analysis is underway to correlate this finding with molecular signatures and potential clinical outcomes.

In 2021 the first relapse appeared with peritoneal dissemination and a tumor recurrence, arising from the rectosigmoid anastomosis site, where the recurring and metastasizing tumor cells were identified, also diffusing into the omental surface. Further, we initiated the analysis of histological images of metastatic MAC tissues, using the Q-Path digital pathology platform (**Figure 5**). Through high-resolution segmentation performed by single-cell annotation, distinct compartments including tumor epithelia, mucinous component, stromal regions, vasculature, and immune cell infiltrates could be identified. This spatial mapping enabled a detailed inventory of cellular and subcellular structures across the tissue architecture. The annotated regions form the basis for integrating morphological features with molecular expression profiles. This integrated analysis is critical for decoding intra-tumoral heterogeneity and pinpointing spatially confined therapeutic targets.

**Figure 5.**
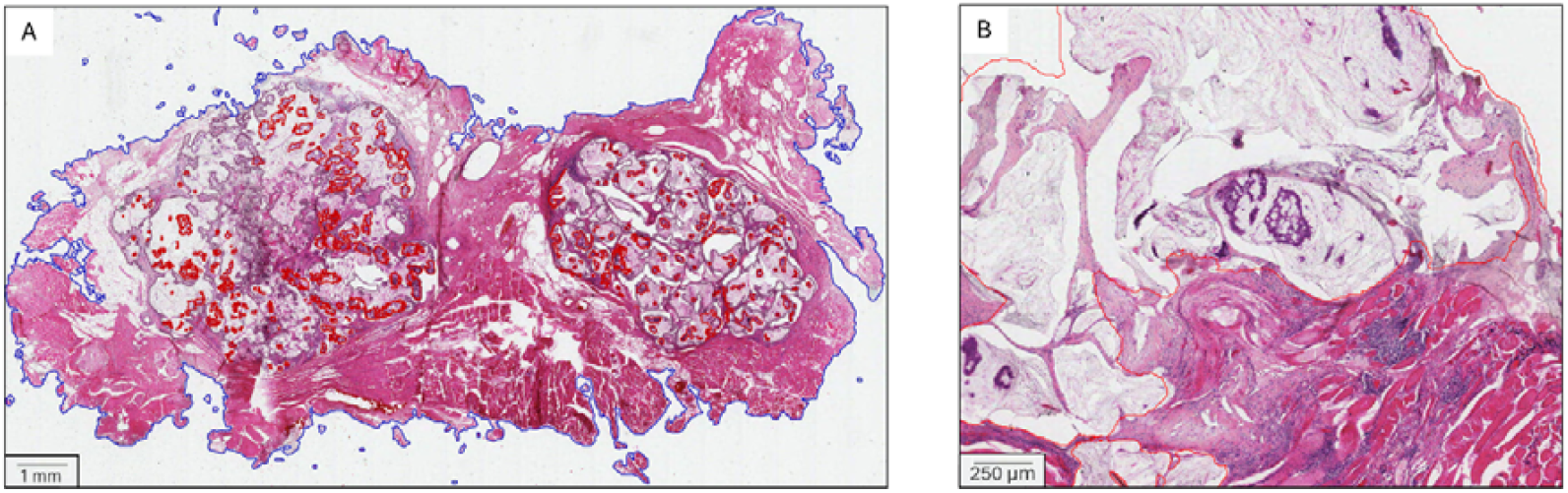
Histological images illustrating the annotated compartments and spatially distinct regions within metastatic MAC in abdominal wall muscle and peritoneal tissues. Using the Q-Path digital pathology platform, we performed systematic segmentation and annotation to identify malignant epithelia, the surrounding mucin, stromal regions, vasculature, and immune cell infiltrates. This workflow enabled a high-resolution inventory of the cellular and subcellular structures within the tissue architecture. The annotations serve as a foundational map for correlating morphological features with molecular expression profiles. Such integration is essential for understanding intra-tumoral heterogeneity and identifying region-specific therapeutic targets. Sample region in **(A)** is with blue outline, and **(B)** with red outline. Tumor tissue is in red. Scale bars: **(A)** 1 mm; **(B)** 250 µm

The re-resected sigmoid tumor specimen, which included regions of recurrent disease at the anastomosis site, was carefully examined to identify and highlight the areas containing recurrent components. These regions were distinctly visualized within the broader tumor architecture, allowing for clear demarcation of metastatic foci and zones of local recurrence. High-resolution imaging and histopathological analysis provided detailed morphological characterization of these recurrent areas, revealing structural differences compared to the primary tumor mass arising from adenomatous polyp. As shown in **Figure 6**, this spatial mapping offers important insights into tumor progression dynamics and recurrence patterns. Such analysis is critical for understanding the biological behavior of residual disease and for guiding future therapeutic strategies.

**Figure 6.**
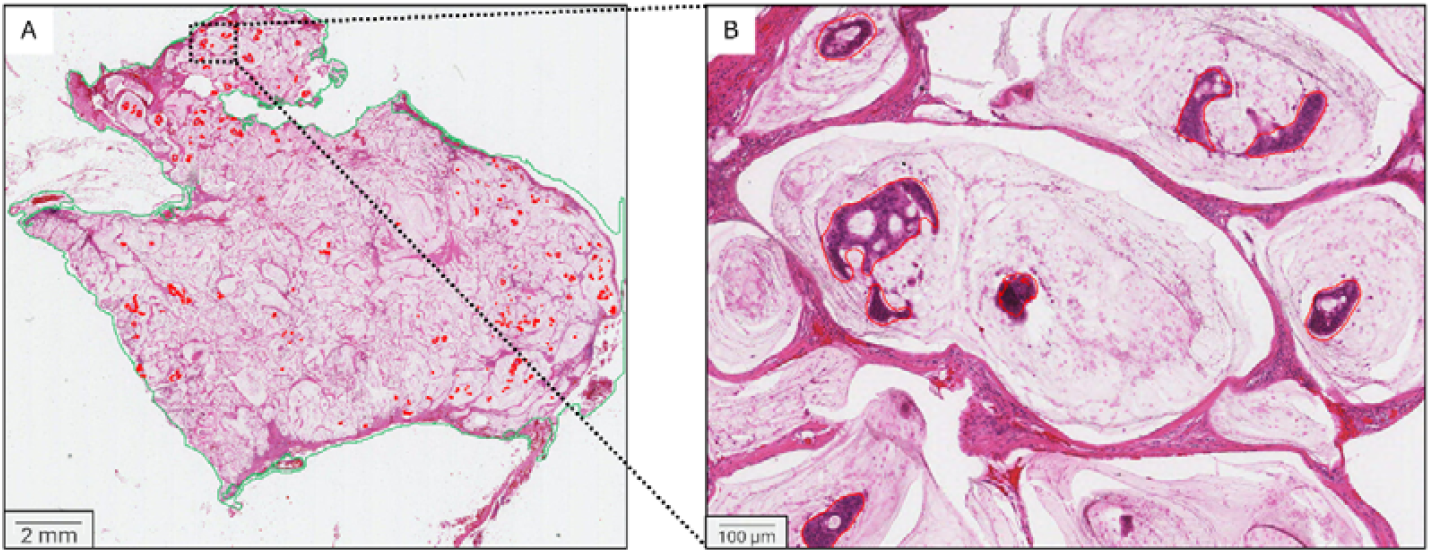
Representative histological images from the re-resected sigmoid tumor arising at the previous anastomosis site, illustrating both metastatic tumor architecture and areas of local recurrence. The sections highlight morphologically distinct regions within the tumor mass, including zones of dedifferentiation and increased mitotic activity characteristic of recurrence. Hematoxylin and eosin staining reveals the infiltrative tumor front adjacent to normal colonic sub-serosal adipose tissue, along with stromal remodeling and inflammatory cell infiltration. These features are consistent with histopathological markers of local tumor regrowth. The annotated images serve as a reference for further molecular and spatial proteomic analyses. Sampled regions are outlined with green color. Tumor cell clusters in red. Scale bars: **(A)** 2 mm; **(B)** 100 µm

The first distant intraabdominal, intraperitoneal metastasis that we observed and that we analyzed originated in 2021, emerging after the completion of chemotherapy treatment. This metastatic spread was followed by additional tumor cell invasions detected on the right side of the diaphragm as well as in the abdominal wall musculature, indicating an increasing burden of disease. In January 2023, the patient was further diagnosed with prostate cancer, representing a significant comorbid malignancy that added complexity to the overall tumor profile. The treatment regimen was adjusted accordingly, and the patient received hormonal therapy (Bicalutamide) in combination with radiotherapy, which successfully stabilized the tumor for a period of approximately two years. However, in May 2023, disease progression resumed with the appearance of peritoneal metastases and tumor spread to the liver ligament, as confirmed by radiological imaging and illustrated in **Figure 7**.

**Figure 7.**
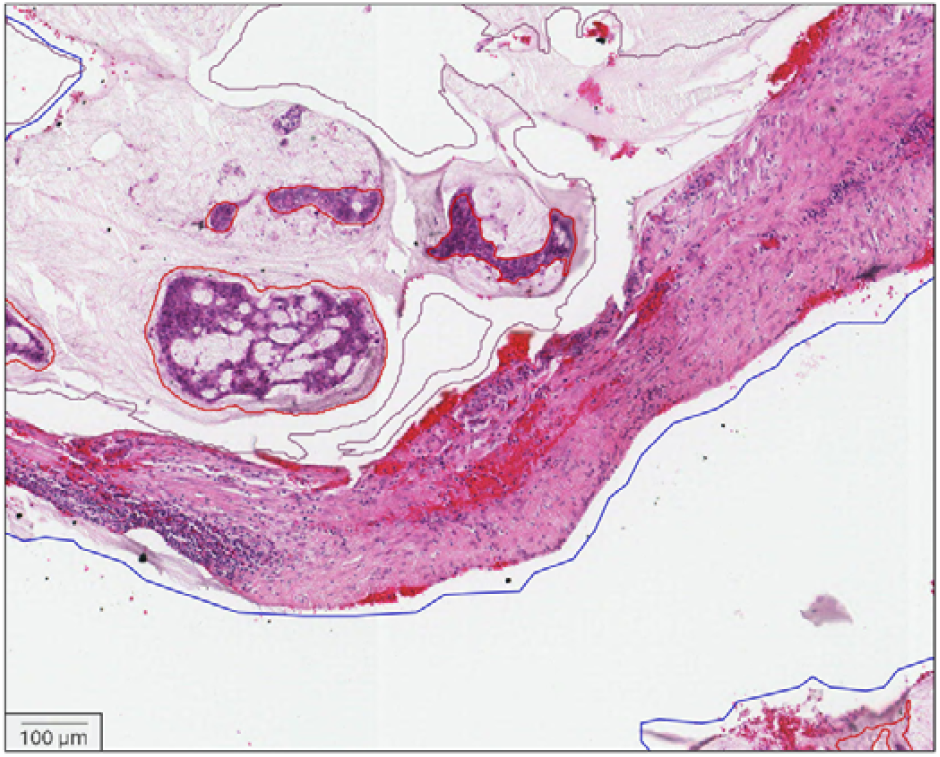
Representative imaging of peritoneal metastasis, showing connective tissue involvement. This excessive amount of extracellular mucin plays an important role of immune system and therapeutic drug resistance, as lymphocytes (shown at the lower left corner) cannot infiltrate this thick barrier. Also notable, that different epithelial manifestations of the tumor are present (such as C-epithelial clumps and layers, D-floating individual tumor cells and, E-complex glandular and cribriform pattern). Sample region is outlined with blue color. Tumor cell clusters or tumor tissue are in red, adipose tissue is in brown. Scale bar: 100 µm

### 3.4. Proteomic findings of peritoneal and Hepatoduodenal metastases

Relative to the primary tumor, the peritoneal metastasis exhibited a marked shift toward an inflamed, stromal–remodeling and invasive program while maintaining an immune-excluded architecture on pathology (inflamed stroma, cold tumor nests). Hallmark/Reactome enrichment showed coordinated upregulation of inflammatory, interferon and NF-κB axes, together with IL-6/JAK/STAT3 and IL-2/STAT5 signaling, alongside TGF-β signaling, EMT and angiogenesis. Complement and coagulation cascades were likewise elevated, consistent with an activated peritoneal microenvironment. Growth and stress-adaptation pathways (mTORC1 signaling, KRAS signaling UP, unfolded protein response) were also positively enriched. In parallel, metabolic tone was reduced, with negative enrichment of fatty-acid metabolism, oxidative phosphorylation and adipogenesis (**Figure 8**).

**Figure 8.**
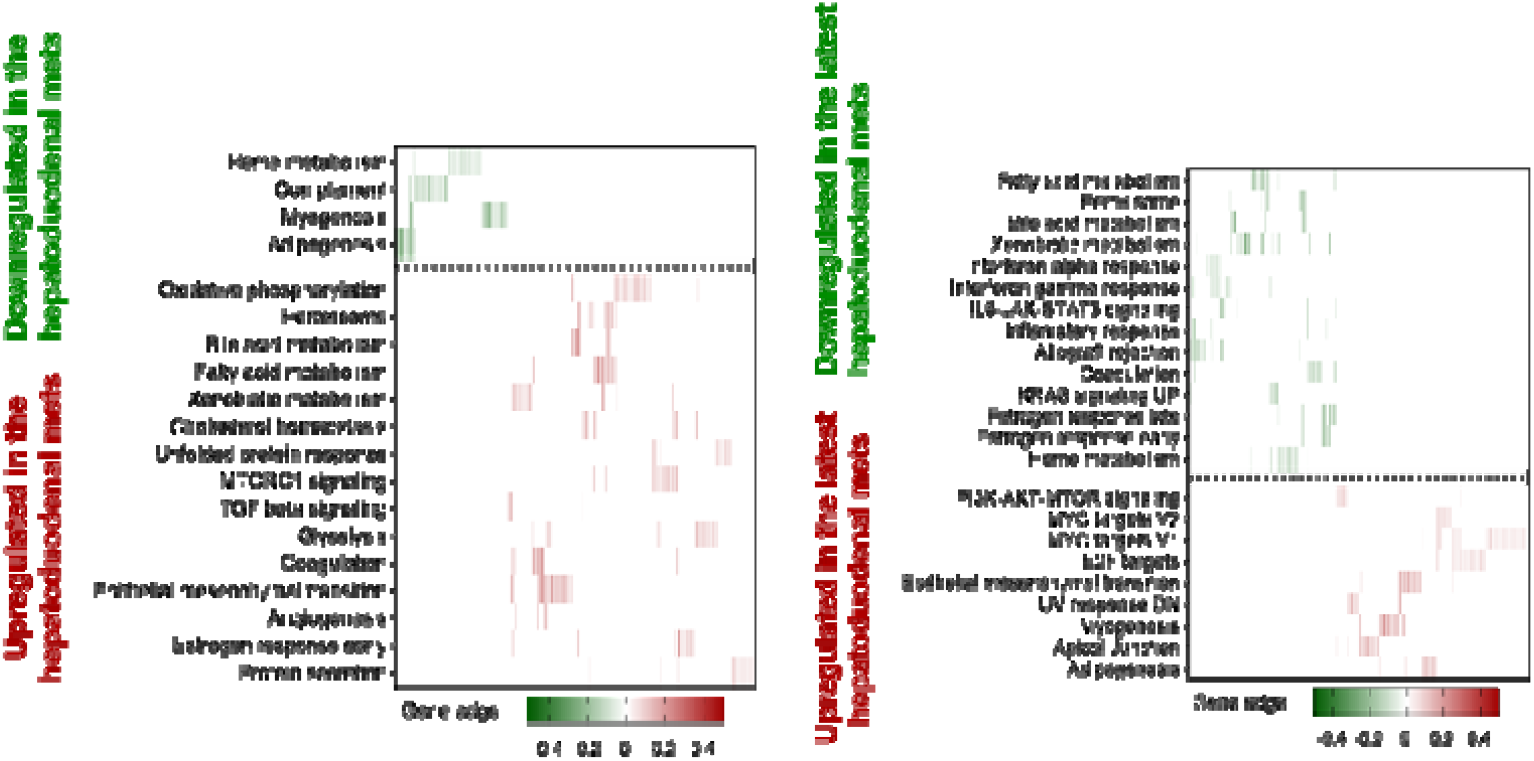
Heatmap showing differential protein regulation in metastatic colontumors. Upregulated proteins are shown in red and downregulated proteins in green, grouped by functional categories. Metastatic tumors exhibit marked upregulation of proteins associated with tumor-promoting pathways.

Consistent with the mucinous phenotype observed histologically, mucin components remained prominent: MUC2 persisted at high abundance, with MUC5AC and MUC4 increased, supporting a continued barrier milieu around tumor cell clusters and aligning with the immune-exclusion pattern noted on digital pathology.

Compared with the peritoneal lesion, the hepatoduodenal metastasis showed a clear shift toward enhanced metabolic activity and plasticity, together with a stress/secretory phenotype and reaffirmation of tissue remodeling pathways. Positive enrichment was observed for oxidative phosphorylation, fatty-acid metabolism, peroxisome, bile-acid metabolism, xenobiotic metabolism, and cholesterol homeostasis, indicating broad lipid and mitochondrial retooling consistent with adaptation to the hepatobiliary niche. Concomitant increases in the unfolded protein response, mTORC1 signaling, protein secretion, and glycolysis support heightened proteostasis demand and anabolic flux. EMT, angiogenesis, TGF-β signaling, and coagulation were also positively enriched, aligning with an aggressive, invasive program and ongoing stromal remodeling. In contrast, heme metabolism, complement, myogenesis, and adipogenesis were negatively enriched. Overall, these changes are consistent with a metastasis that is more metabolically active and stress-adapted, with molecular features compatible with greater dissemination potential (**Figure 8**).

Relative to the immediately preceding metastasis, the latest lesion displayed further metabolic/inflammatory reprogramming, with positive enrichment of estrogen response (early/late), KRAS signaling (UP), IL-6/JAK/STAT3, interferon-α/γ responses, inflammatory response, allograft rejection, and coagulation. Metabolic pathways were again positively enriched. In parallel, there was negative enrichment of E2F targets, MYC targets (V1/V2), PI3K–AKT–mTOR signaling, epithelial–mesenchymal transition, apical junction, UV response DN, adipogenesis, myogenesis, and heme metabolism (**Figure 8**). Taken together, these data indicate a transition away from a cell-cycle/MYC-dominant profile toward an inflammation-centered, lipid/oxidative metabolism-oriented state, while maintaining features supportive of tumor–stroma interaction (coagulation/inflammation).

### 3.5. Colon Mucinous Barrier Formations Within Primary and Metastatic Tumor Tissue

Mucinous adenocarcinomas of the colon are characterized by abundant extracellular mucin production, which forms a distinct barrier within the tumor microenvironment. This mucinous barrier, composed primarily of MUC2 and related glycoproteins, is secreted by neoplastic epithelial cells and constitutes more than 50% of the tumor volume in classic mucinous subtypes. Within the primary tumor, this barrier can physically separate cancer cells from immune infiltrates and therapeutic agents, contributing to immune evasion and chemoresistance. Additionally, the mucin matrix alters the mechanical properties of the tumor, facilitating local invasion and dissemination through the disruption of epithelial integrity and promotion of tumor budding. In June 2024, additional surgery was undertaken for the aim of removing tumor burden caused by the peritoneal metastases, that this time also included sub-cutaneous dissemination, and furthermore, abdominal wall metastases as well. At this disease-stage, upon histopathological examination, the presence of the complex glandular/Cribriform phenotype increased greatly. We were unable to microscopically detect any lymphocyte infiltration within the tumor regions. This absence may be due to the mucinous barrier, which effectively prevents immune cell entry. Notably, lymphocytes were observed only in areas beyond the mucin-rich zones that surround and protect the five identified tumor phenotypes (see **Fig. 9**).

**Figure 9.**
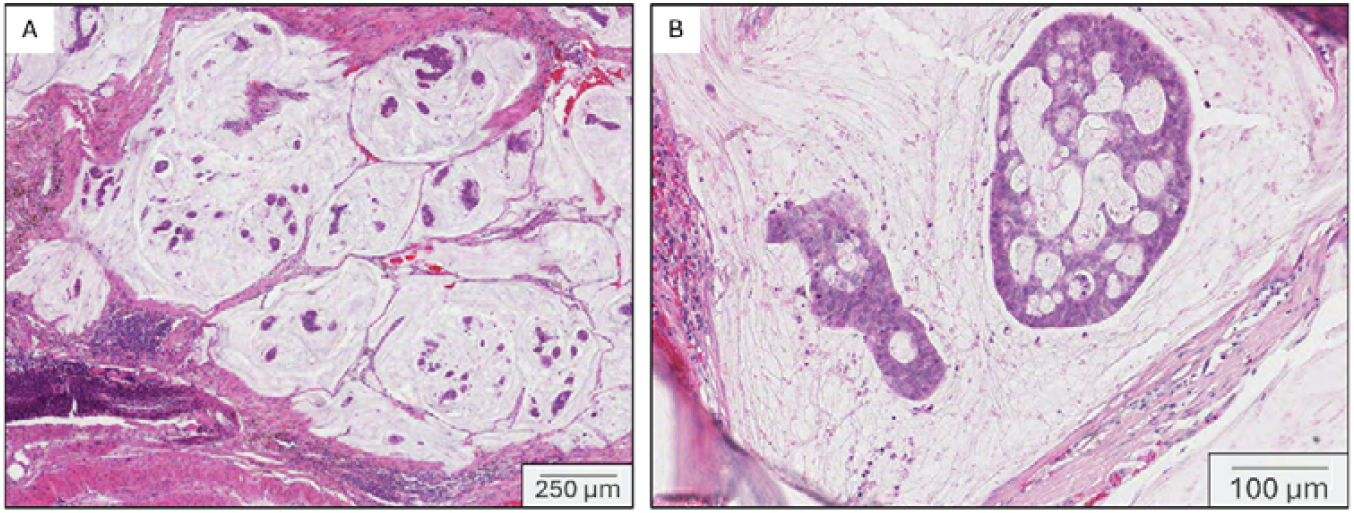
(A) Representative imaging of peritoneal metastasis, showing connective tissue involvement. This excessive amount of extracellular mucin plays an important role of the immune system and therapeutic drug resistance, as lymphocytes (shown at the lower left corner) cannot infiltrate this thick barrier. (B) Cribriform/complex glandular morphology of tumor epithelia appeared mostly in late metastatic samples. Scale bars: **(A)** 250 µm; **(B)** 100 µm

Machine based AI imaging and digital pathology analysis demonstrate that this mucinous colorectal cancer has progressed from a polyp to peritoneal dissemination with abdominal wall invasion despite repeated cytoreductive surgery and HIPEC. Spatial and molecular profiling suggests that the heterogeneous mucin architecture ranging from dense barrier-like deposits to dispersed mucus functions not only as a physical impediment to drug delivery but also as an active modulator of the tumor immune microenvironment, directly influencing disease progression and therapeutic response.

## 4. DISCUSSION AND CONCLUSIONS

The combined analysis of mucin production and tumor proteomics with histopathology reveals crucial mechanistic insights with significant therapeutic implications. A prominent feature in the tumor microenvironment is the strong mucin barrier, particularly involving MUC2, which forms a dense extracellular matrix that impedes immune cell infiltration and hinders drug penetration. The tumor contains areas where dense, barrier-like mucin coexists with softer, more diffusely distributed mucus. Both histological assessment and advanced molecular analyses, including proteomic profiling, demonstrate that mucin functions not only as a physical barrier to drug penetration but also as an active modulator of immune responses and the tumor microenvironment. This aligns with previously published findings (19) in the management of low-grade mucinous neoplasms and pseudomyxoma peritonei (PMP). In such contexts, mucolytic agents, most notably the combination of bromelain and N-acetylcysteine, have demonstrated promising mucolytic activity, facilitating improved delivery of therapeutic agents (20–24). In PMP, intraperitoneal administration of mucolytics has shown promise in softening mucinous deposits and enhancing treatment efficacy, offering a rationale to explore similar strategies in mucin-rich malignancies (20,21,24).

The barrier limits immune surveillance, modifies the metastatic microenvironment, and sustains proliferative signaling through the sequestration of cytokines and growth factors (25). Recent proteomic and transcriptomic profiling has revealed that the mucin-producing phenotype is maintained through distinct gene expression programs, often involving KRAS mutations (26). Understanding the structural and biochemical properties of mucinous barriers in both primary and metastatic settings offers promising avenues for targeted therapy, including mucolytic agents and drug delivery systems designed to penetrate these protective matrices (24). Complementary molecular profiling data provides a layered understanding of tumor progression and heterogeneity. Proteomic comparisons between the primary tumor and its metastases indicate a shift in biological behavior and microenvironmental features. The primary tumor is characterized by genomic instability and heightened proliferative signaling, with limited metabolic and inflammatory activity, potentially due to low vascularization and heavy mucin content. Peritoneal metastases, in contrast, show marked immune modulation, inflammatory signaling, epithelial-to-mesenchymal transition (EMT), and early angiogenesis, indicative of adaptation to a new niche. Hepatoduodenal metastases exhibit even greater complexity, with increased metabolic flexibility, advanced EMT signatures, and activation of aggressive signaling axes such as PI3K/AKT/mTOR and MYC.

These findings underscore the need for a multi-faceted therapeutic approach. Beyond enzymatic mucin degradation to improve drug accessibility, treatment strategies must also address proliferative capacity, immune evasion, and metabolic reprogramming. Together, these insights suggest that personalized therapeutic regimens incorporating mucolytics, targeted inhibitors, and immunomodulatory agents may offer superior outcomes in mucin-producing and metastatic malignancies.

The proposed integrated treatment strategy addresses the complex challenges posed by mucin-rich metastatic tumors through a multifaceted approach, combining mucolytic, targeted molecular, immunologic, and microenvironment-directed therapies. Initiating or continuing mucolytic therapy with a combination of bromelain and N-acetylcysteine (BromAc) may help dismantle the physical barrier imposed by dense mucin, thereby enhancing therapeutic penetration and reducing tumor-protective stromal effects. Given bromelain’s favorable safety profile, both systemic and intraperitoneal delivery are feasible and warrant further exploration in clinical settings (27). The reduction of mucin burden may potentiate the efficacy of concurrent molecular therapies.

Targeted inhibition of proliferative signaling pathways is a logical next step. The presence of elevated E2F and G2/M checkpoint signatures in metastatic samples supports the inclusion of CDK4/6 inhibitors to suppress cell cycle progression (28). Furthermore, hepatoduodenal metastases showing strong MYC and PI3K/AKT/mTOR activation suggest potential sensitivity to mTOR inhibitors or emerging MYC-directed therapeutics (29). The immune-active microenvironment observed in peritoneal metastases introduces opportunities for immunotherapy, including immune checkpoint blockade and IL6/JAK/STAT3 pathway inhibition (30,31). Additionally, considering the role of retinoid signaling in mucin synthesis, targeting retinoic acid receptors may further reduce mucin production and amplify the impact of mucolytics (32,33).

Addressing the tumor microenvironment remains critical for sustained disease control. Angiogenic activity in metastatic compartments is hardly present morphologically in our case, however, the use of anti-VEGF agents like bevacizumab can be considered to limit neovascularization (34). Depending on molecular re-assessment, metabolic modulators such as metformin or phenformin may be integrated to counteract metabolic adaptation in colon tumors (35). Continued surveillance through imaging modalities (MRI, FDG-PET) and biomarker monitoring (e.g., CEA) will guide dynamic adjustments to the therapeutic regimen (36). Incorporating re-biopsy and molecular re-profiling will further refine treatment decisions. Finally, the implementation of this strategy requires close multidisciplinary coordination across oncology, pathology, surgery, and interventional specialties. This integrated model not only reflects current precision oncology paradigms but also aims to disrupt the protective tumor-stroma interactions that fuel progression, thereby improving long-term clinical outcomes.

## Patient consent statement

Written informed consent was obtained from the patient for participation in the study and for publication of clinical data and associated images.

## Author Contributions

L.F., B.S., J.G., and G.M.V contributed equally to this work. L.F. and B.S. designed and performed histopathology analysis. J.G., N.W., and J.G. performed experiments and analyzed data. L.F., B.S., J.G., G.M.V and L.S., J.M., I.B.N. and M.A. S. wrote and edited the manuscript. L.C., L.S., L.F. and B.S. contributed to visualization and validation. J.G., M.M.V., R.A., K.P., L.S., and L.C. contributed to the study design, conducted histopathology analysis, and assisted with data analysis. L.S., J.G., E.W., K.P., J.M., I.B.N., M.A.S., G.M.V edited and supervised the manuscript.

## Acknowledgements

The authors used ChatGPT (OpenAI, GPT-5.1)(37) to assist with language refinement and improve readability during manuscript preparation. All AI-assisted content was subsequently reviewed and edited by the authors, who take full responsibility for the final version of the publication.

## Data Availability Statement

Proteomics data generated in this study was deposited to the ProteomeXchange Consortium via the PRIDE partner repository. The accession number will be provided during submission.

